# An ultrasound absorbing inflorescence zone enhances echo-acoustic contrast of bat-pollinated cactus flowers

**DOI:** 10.1101/2019.12.28.890046

**Authors:** Ralph Simon, Felix Matt, Vinicio Santillan, Marco Tschapka, Merlin Tuttle, Wouter Halfwerk

## Abstract

Flowering plants have evolved an extraordinary variety of signaling traits to attract and guide their pollinators. Most flowers rely on visual and chemical signals, but some bat-pollinated plants have evolved reflective surfaces to acoustically guide echolocating bats. All known acoustic flower signals rely on the same principle of increased sonar reflectivity. Here we describe a novel mechanism through which plants can make flowers acoustically conspicuous, a principle that relies on increased absorption of the area surrounding the flower. In a bat-pollinated columnar cactus (*Espostoa frutescens*) we found a hairy inflorescence zone, a so called cephalium. Flowers solely emerge out of this zone. We measured the ultrasound echoes of cephalia, flowers and unspecialized column surfaces and recorded echolocation calls of approaching bats. We found that the cephalium acts as strong ultrasound absorber, attenuating the sound by −14 dB compared to other parts of the column. The absorption was highest around the echolocation call frequencies of approaching bats. Our results indicate that, instead of making flowers more reflective, plants can also evolve structures to attenuate the background echo, thereby enhancing the acoustic contrast with the target. Similar sound absorbing mechanisms may be found in other species that interact with bats across a wide range of ecological contexts.

## Introduction

Flowering plants rely on a wide variety of communication strategies to attract their pollinators. Conspicuous visual flower signals are in particular useful to guide receivers, as they are easy to locate and the use of colours makes flowers stand out against the vegetation background (Spaethe *et al*. 2001; Forrest & Thomson 2009; Trunschke *et al*. 2021). Neotropical bat-pollinated plants are limited in the use of visual signals to attract their pollinators and therefore independently evolved acoustic traits (Simon *et al*. 2021) or even echo-reflective structures to acoustically guide these nocturnal pollinators (von Helversen & von Helversen 1999; von Helversen *et al*. 2003; Simon *et al*. 2011). Echo-acoustic signalling plants all use concave shapes with either triple mirror, bell- or dish-like structures. These concave shaped structures share the same basic acoustic principle of focusing returning echoes to an approaching bat, thereby increasing the range over which they can be detected. Some flower signals use additional spectral-temporal signatures increasing conspicuousness (Simon *et al*. 2011). These echo signatures are generated by interferences between sound pathways on the reflector surface that cause enhancement of certain frequency bands (Simon *et al*. 2020). Reflective plant structures not only evolved in the Neotropics but also in a bat dependent pitcher plant from Borneo. The pitcher plant *Nepenthes hemsleyana*, depends on bats roosting inside the pitcher as they provide additional nitrogen intake through their droppings (Grafe *et al*. 2011). These plants therefore developed a reflective prolonged pitcher backwall to advertise their pitcher-leaves as roosts (Schöner *et al*. 2015). The fact that reflective plant structures independently evolved several times in different ecological contexts and in different plant families shows how importat such signals are for ecological networks.

Here we assess an evolutionary novel adaptation that enhances acoustic communication between plants and pollinating bats. Some cacti species exhibit inflorescence zones on their column that are particularly hairy, the so-called *cephalium* (see Fig. S1). There are several different morphologies of cacti described as cephalia, and we refer here to what is described as a lateral cephalium by Mauseth (2006). Several functions of these *cephalia* zones, have been proposed. The hairy structure may shield buds from UV radiation at high altitudes, or protect against nectar robbers and herbviores (Buxbaum 1961; Martorell *et al*. 2006; Mauseth 2006). Here we test a hypothesis by von Helversen et al. (2003), which states that such hairy zones may have been co-opted to serve in bat-pollinated cacti as sound-absorbing structures that support detection and localization of sound-reflecting flowers by pollinating bats.

Using a bat-mimetic sonar-head we carried out ensonification experiments with different structures of the cactus *Espostoa frutescens* (von Helversen *et al*. 2003) from the Andes. Specifically, we ensonified the cactus’ column, flowers as well as the hairy *cephalium* zone. Additionally, we recorded the echolocation calls of its main pollinator, the nectar-feeding bat *Anoura geoffroyi* (Phyllostomidae) and assessed whether the cephalium was especially absorbent in the ultrasonic frequency range of the calls.

## Material and Methods

We studied *Espostoa frutescens* and its pollinator, Geoffroy’s tailless bat (*Anoura geoffroyi*). The study was carried out in a dry valley of the Ecuadorian Andes, close to the city Oña in the province of Azuay. As it was not possible to conduct the echo measurements in the field – the cacti are growing in rocky and steep habitat - we had to cut some columns (n = 6) and conducted the measurements indoors at a nearby farm. All experiments where approved by the local authorities (Ministeria del Ambiente, Cuenca, Ecuador, autorizacion para investigación científica N. 035-DPA-MA-2012). A specimen of *Espostoa frutescens* is deposited at Herbario Azuay (Cuenca, Ecuador) with the number HA 7814.

To measure the reflectance of the different structures of the cacti we mounted the columns on tripods and used a custom-built biomimetic sonar-head to ensonify them. The sonar-head consisted of a 1/4” condenser microphone (40BF; preamplifier 26AB; power module 12AA; G.R.A.S. Sound & Vibration, Holte, Denmark) and a custom-made EMFi (Electro Mechanical Film) loudspeaker (sound pressure levels at 1 m distance: 92 dB ± 8 dB, frequency range: 30-160 kHz; Department of Sensor Technology, University of Erlangen-Nuremberg, Erlangen, Germany). The speaker and the microphone were embedded in an aluminium body and were placed next to each other, similar to how the mouth/nose and ears are arranged on a bat’s head. To ensure a quite narrow sound beam similar to the one of a nectar feeding bat, we ensonified cacti from a relatively short distance of 15 cm. As we measured the impulse responses (echo responses of very short pulses) we had no problem with overlap. We obtained the impulses by ensonifying with a continuously replayed MLS Signal (Maximum Length Sequence a pseudorandom noise like signal) and recorded the reflected sound. A basic property of any MLS signal is that their autocorrelation function is perfectly narrow, therefore it is possible to obtain the impulse responses (IRs) by deconvolution of the reflected echo and the original MLS (von Helversen *et al*. 2003). The advantage of ensonifying with an MLS instead of any bat like sweep or short impulse is that it is possible to ensonify with more sound energy and therefore the obtained echoes have a much better signal to noise ratio. The spectra of the echoes were obtained by windowing the IRs (rectangular, 1024 samples) around the echo of the cactus and then calculating the power spectral density (PSD). To obtain spectral target strength (TS), independent of the frequency response of the loudspeaker, we conducted another measurement. For this measurement the cactus was replaced by an acrylic glass plate oriented perpendicular to the direction of sound propagation. We also deduced PSD of the total reflection of that acrylic plate and then calculated the difference between PSD from the recordings of the cactus surfaces and the PSD of the “total reflection” of the acrylic glass plate (For more information on the setup and the signal processing see also (von Helversen *et al*. 2003; Simon *et al*. 2006; Simon *et al*. 2011)).

Using our ensonification setup we measured the acoustic properties of six freshly cut columns of *E. frutescens*, focusing on the hairy cephalium zone and the unspecialized parts (backside) of the column. For both measurements we scanned the columns by moving the sonar-head upwards along its vertical axis and made 10 measurements at different heights of the column. We also measured the reflectance of six isolated flowers, which were mounted on a stepping motor. We rotated the flower in 6° steps and measured 10 echoes around the opening (0°) of the flowers from −30° to 30°.

To deduce an average target strength for each structure and each cactus we averaged the target strength for the 10 recordings we had of every structure (see S2). We also averaged TS for 5 different frequency bands. One frequency band covered the whole bandwidth which we could measure with our setup. The other frequency bands were chosen to split the whole bandwidth in four logarithmically spaced frequency bands. We did this because auditory perception shows an exponential frequency distribution along the cochlea, see (Simon *et al*. 2006) for more information on spectral echo perception.

To understand how the echo of a flower would be received if *E. frutescens* had no cephalium structures we manipulated one column and mimicked a flower which grows on an unspecialized part of the column. We first scanned the hairy cephalium with an open flower by moving the sonar-head upwards along the vertical axis of the column over an area of 30 cm. The flower was located central on this area and we measured in 1 cm steps. After the measurements we resected the flower from the cephalium and fixed it on the hairless backside of the column (Fig. 2B). For this experimentally manipulated column we made the same detailed vertical scan (30 cm, 1 cm steps).

**Fig. 1.**
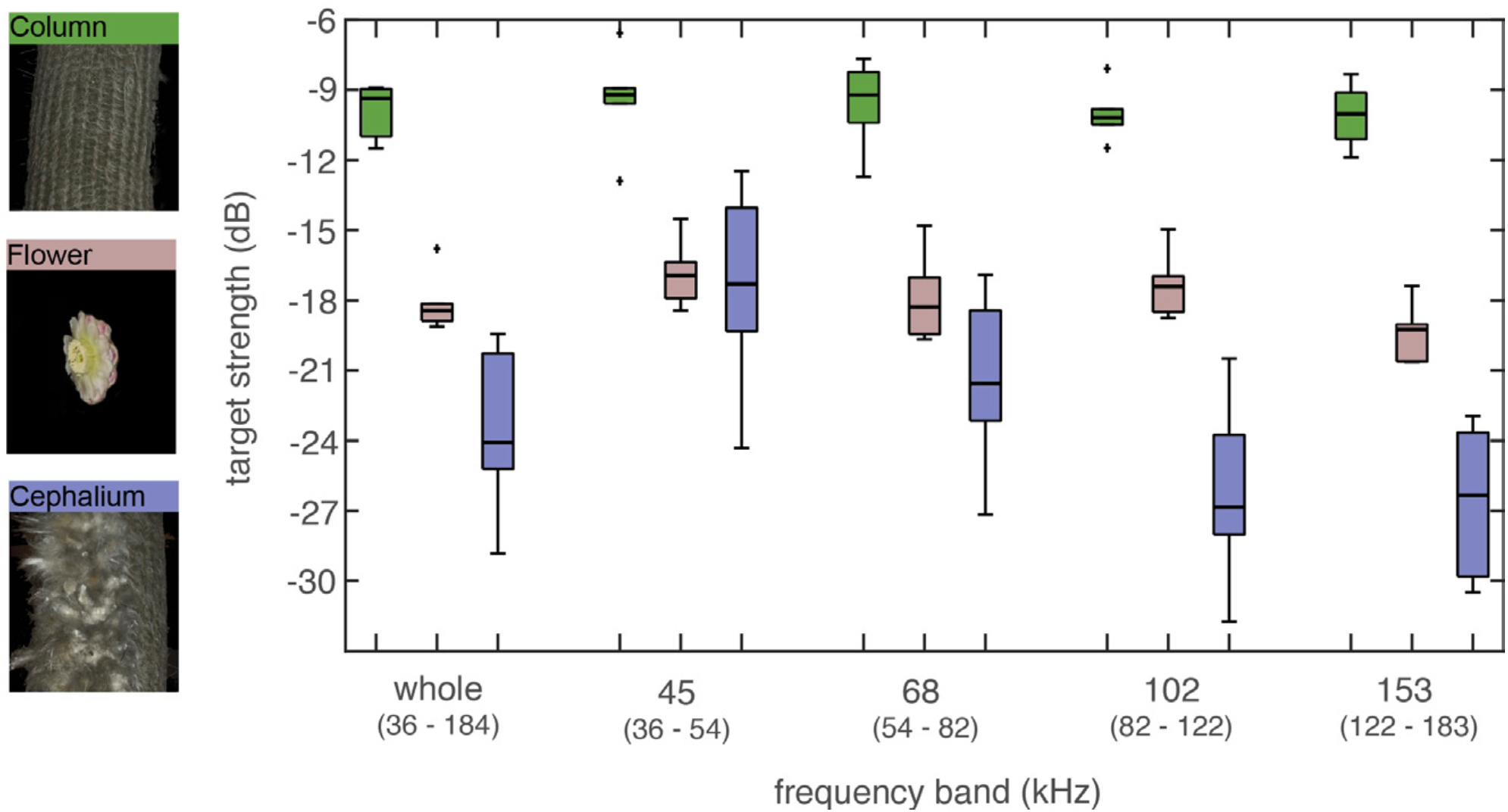
Spectral target strength (TS) of different morphological structures of *Espostoa frutescens* for different frequency bands. The spectral target strength was obtained from ensonification measurements at a distance of 15cm. We measured unspecialized parts of the cactus column (green boxplots; n = 6 columns, 10 measurements per column), isolated flowers (rose boxplots; n = 6 columns, 10 measurements per flower from different angles) and the hairy cephalium zone (purple boxplots; n = 6 columns, 10 measurements per column).

**Fig. 2.**
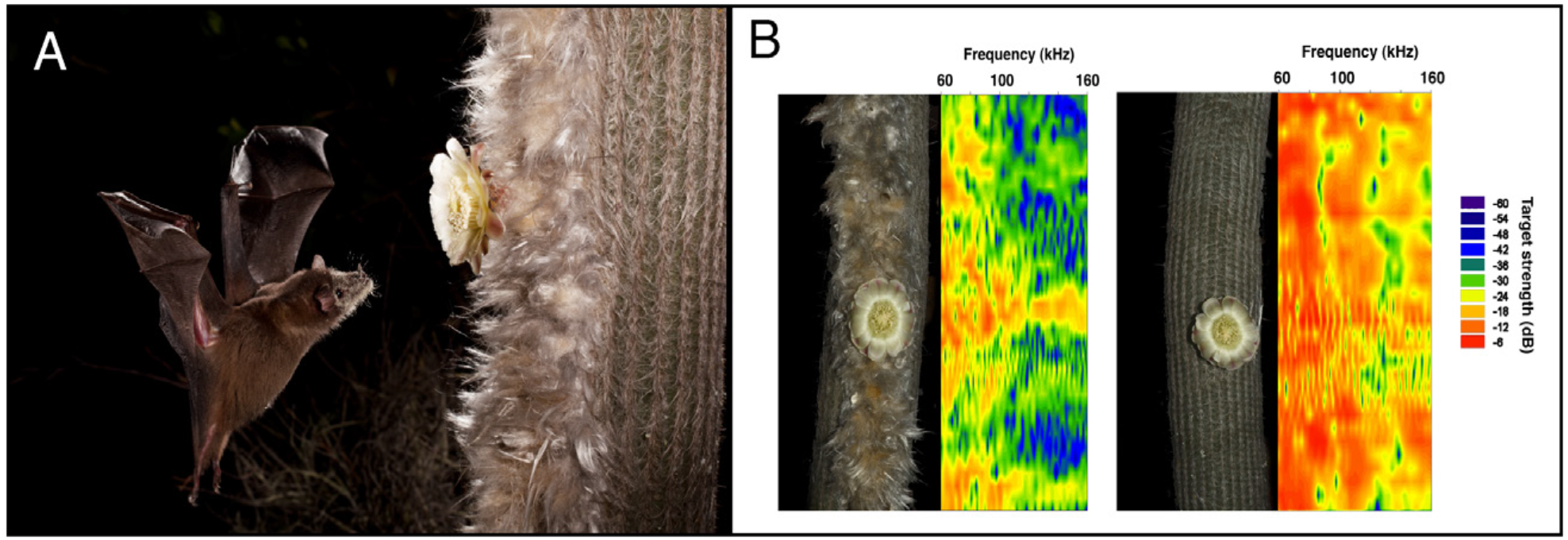
Nectar-feeding bat approaching a flower and echo fingerprints of different cactus surfaces with flowers. (**A**) Image of a Geoffroy’s tailless bat (*Anoura geoffroyi*) approaching a flower of *Espostoa frutescens*, which is in embedded in the hairy cephalium zone (photo credit: Merlin Tuttle’s Bat Conservation). (**B**) Echo fingerprints of acoustic scans along the cactus column. The left column is a natural column with cephalium and flower, for the right measurement we experimentally manipulated the column. The flower was cut out of the hairy zone and fixed on an unspecialized part of the column. The intensity (spectral target strength in dB) of the echo is given in colour gradation (red indicates high intensities, blue low intensities).

We also recorded echolocation calls of two male Geoffroy’s tailless bats (*Anoura geoffroyi*)approaching an *Espostoa* column with an open flower. The microphone (1/4” condenser microphone 40BF; preamplifier 26AB; power module 12AA; G.R.A.S. Sound & Vibration, Holte, Denmark) was placed next to the flower (see Fig. S3) and we recorded with a sampling rate of 500 ks/s. We obtained 45 manually triggered recordings, each with a length of 2 s, during the approaches of the bats. To ensure a good signal-to-noise-ratio for the call analysis we selected 21 approach sequences where at least two calls had a max amplitude of more than 60 mV. We analysed the calls using the program Avisoft-SASLab Pro (Avisoft Bioacoustics, Glienicke, Germany).

We tested for significant effects of plant structure on echo-acoustic target strength using the lmer package in R (version 3.5.3). We constructed linear mixed models and checked model assumptions by visual inspection of the residuals. Target strength was averaged over the 10 measurements per plant individual and structure and modelled as dependent variable. Plant structure (column, flower or cephalium) was added as fixed factor and plant individual as a random intercept term. For the different frequency ranges, we modelled the interaction between plant part and frequency band. We tested for significance of main effect of plant structure on target strength and for significance of the interaction between structure and frequency band by comparing models with and without terms using likelihood ratio tests.

## Results

We found a significant effect of plant structure on overall target strength (LMM, n = 18 plant structures, n = 180 measurements, d.f. = 2, X2 = 39.31, P < 0.001). Furthermore, target strength depended on the interaction between frequency range and plant structure (LMM, d.f. = 8, X2 = 37.51, P < 0.001). Overall, the plain column surface of *E. frutescens* reflected the strongest echoes. We measured a high target strength (average TS −9.8 dB) for these unspecialized surfaces of the cactus across a wide range of frequencies (Fig. 1). The overall average TS of the flower was much lower compared to the column (−18.1 dB) but also remained similar across all measured frequency bands (Fig 1). The hairy cephalium zone on the other hand showed differences in TS for the different frequency bands (Fig 1). For the lower frequency band (45 kHz) the TS was about the same level as the flower (−17.5 dB) but for higher frequency bands it was much lower, down to −26.3 dB for the 102 kHz frequency band. Overall, the cephalium zone had an average target strength of −23.7 dB, which is around 14 dB lower than the unspecialized parts of the column. The averaged full spectra (Fig. S2) of the different structures showed the same pattern. Flower spectra and cephalium spectra start to diverge from 70 kHz up to higher frequencies with a maximum at around 150 kHz, where they had an average difference of over 10 dB.

A qualitative analysis of the echo-acoustic fingerprint of specialized versus unspecialized parts of the column revealed more detailed insight into the effect of the background on detectability of flower targets (Fig 2B). The unspecialized column reflects high TS echoes for almost the entire bandwidth, which are only sometimes interrupted by some frequency notches. The specialized cephalium side of the column reflects much less sound energy, especially for frequencies above 90 kHz. When scanning the column at the position of the flower, the unmanipulated flower stands out from the less-reflecting background, in particular at frequencies above 90 kHz. When we placed a flower on the unspecialized part of the column the flower echoes almost completely disappear within the loud background echoes, although there might be some additional interference patterns affecting the TS.

In total we analysed 279 echolocation calls of two individuals of *A. geoffroyi*, see Fig. 3 for an example of an echolocation call sequence during an approach to an *E. frutescens* flower. The calls where short, having a duration of only 0.47 ms ± 0.18 ms (mean ± SD) and they were step frequency modulated starting at 132.7 kHz ± 8.1 kHz and ending at 59.8 kHz ± 10.0 kHz. The peak frequency of the calls was at 92.5 kHz ± 4.4 kHz, which falls into the frequency band where sound absorption of the cephalium was highest.

**Fig. 3.**
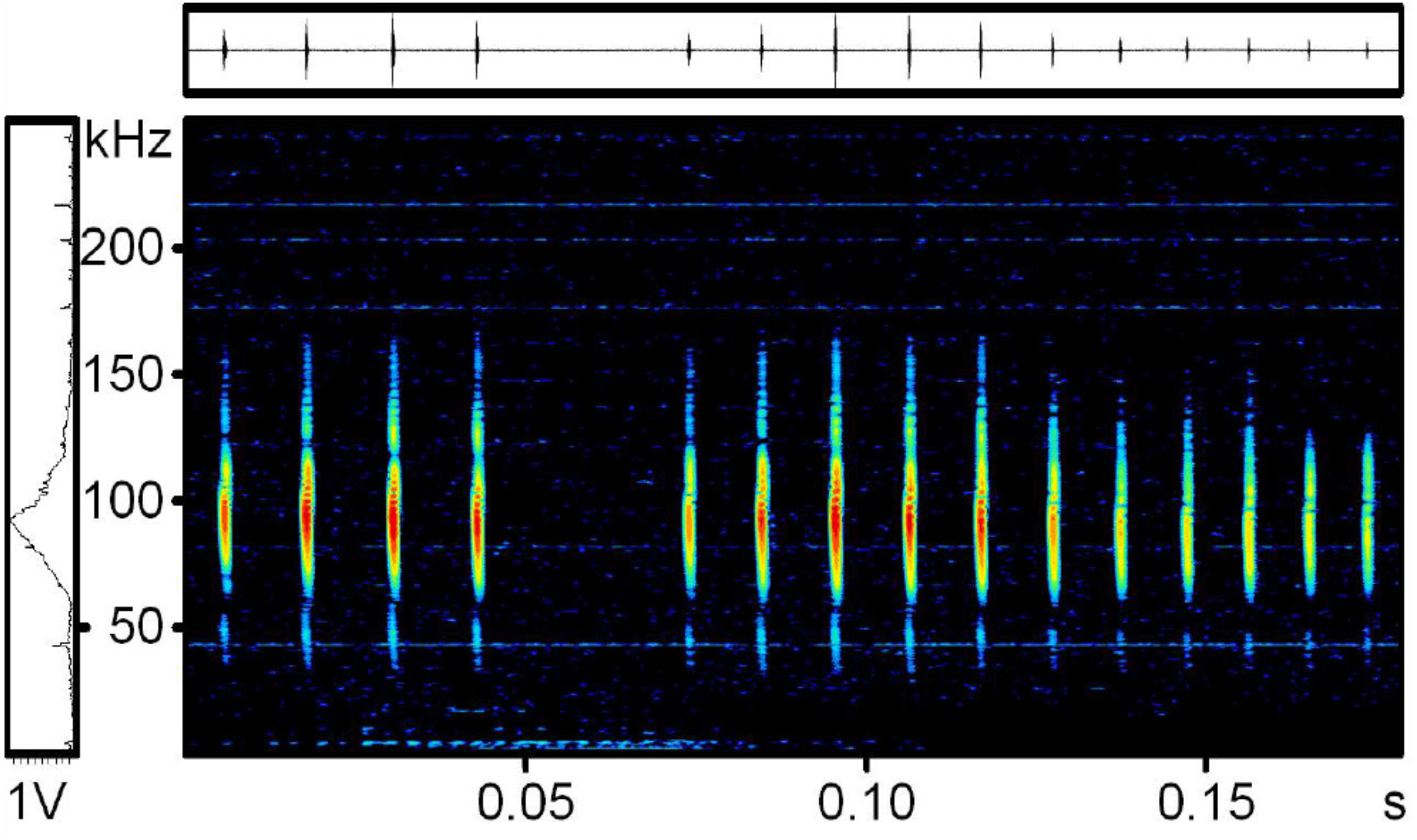
Typical series of calls of a Geoffroy’s tailless bat (*Anoura geoffroyi*) while approaching an *Espostoa frutescens* flower. The microphone (1/4” G.R.A.S. free field microphone) was placed next to the flower as can be seen in Fig. S3.

## Discussion

Our ensonification experiments revealed distinct and frequency-dependent differences in echo-acoustic reflectance of different cacti parts. We found that the plain column of *Espostoa* acts as a strong reflecting surface as it is cylindric, providing reflective surfaces from all directions, and also because the surface has ridges, which may additionally act as small retroreflectors. The flowers of *Espostoa* reflect much less energy compared to the column, mainly due to the fact that the reflecting surface is smaller and that the flowers have a lot of anthers, which scatter sound energy. The specialized cephalium surrounding the flowers reflected the least energy, in particular in the echolocation call frequency range of the plant’s main pollinator, *A. geoffroyi*.These results strongly suggest that the cephalium of *Espostoa* functions as a sound-absorbing structure and thus enhances the echo-acoustic contrast between the flower and the vegetative part of the plant for an approaching bat. While scanning cacti columns for flowers along the cephalium the bats will receive faint echoes unless their call hits a flower, which increases the echo response by around 10 dB. In contrast, flowers growing on the unspecialized parts of the column would be much more difficult to recognize in front of the highly reflective background. Bats might be able to pick up on the interference patterns caused by the flowers, however, this would require much more processing than a salient flower echo in front of an absorbing surface.

As nectar-feeding bats have to visit or revisit several hundred flowers each night to cover their nightly energy expenditure (Winter & von Helversen 2005), this simple yet efficient mechanism of dampening the background of the flowers may help the bats to save on foraging time and thus increase foraging efficiency. The plant on the other hand will benefit from a higher cross pollination rate. Bats are very efficient pollinators that carry a lot of pollen in their fur (see Fig 2A) and have a huge home range so they can pollinate plants growing far apart (von Helversen & Winter 2003).

The absorption of the *cephalium* is most efficient for the 102 kHz frequency band (82 kHz – 122 kHz), which translates to a wavelength of around 3.4 mm (4.2 mm – 2.8 mm). The microstructure of the *cephalium* apparently favours absorption of sound around this wavelength, while larger wavelengths (e.g., 7.6 mm for the 45 kHz band) are around 10 dB less attenuated. The hairs are much smaller in diameter than the wavelengths of sound they absorb best and therefore probably do not scatter the incoming sound waves. An alternative explanation could be that the hairs create a layer of air with different temperature and humidity that reflects the sounds in a frequency-dependent manner.

As other species of *Espostoa* show the same hairy cephalium zone this floral acoustic adaptation might not only be limited to this species and even other genera have similar hairy cephalium zones e.g. *Microanthocereus* (Mauseth 2006). Interestingly, bird pollinated species of the genus *Microanthocereus* have also cephalium zones, however the fur is less dense (personal observation). We argue that cephalium-like structures originally evolved for protection of floral structures, but was co-opted at some point in time to serve an additional or new functional role in pollinator attraction. Once co-opted, the cephalium of bat-pollinated flowers got optimized for this new function through selection by the echolocating bat pollinators.

Our study shows that bat-pollinated flowers can also rely on absorption in addition to reflectance as an acoustic adaptation towards their pollinators. Echoacoustic absorption likely plays a much larger role across a wide range of ecological contexts than so far has been appreciated. Sound absorbent structures have already been described for moth scales (Shen *et al*. 2018) as well as for thoracic moth fur (Neil *et al*. 2018). Whether absorption has adapted in the context of predator-prey arms races remains however to be tested, ideally in a comparative phylogenetic framework.

## Acknowledgements

We are very grateful to Otto von Helversen who initiated this study before his all to early death. We greatly acknowledge the help from Nery Fabian Chamorro Rodas during collection and the measurements of the cacti and the help from all people from Susudel Granja Organica. We thank the Herbario Azuay (Patent: FLR.S-004-2019) for the identification and assistance with the *E. frutescens* specimen.

## Data deposition

All data described in this paper is publicly available at https://github.com/GitRaSimon/SoundAbsorbCephal

## Author contributions

Conceptualization: R.S., F.M., W.H.; Methodology: R.S., F.M., V.S., W.H.; Software: R.S.; Validation: R.S., W.H.; Formal analysis: R.S., W.H.; Investigation: R.S., F.M., V.S., M.Tu. Resources: R.S., F.M., V.S., M.Tu.; Data curation: R.S., W.H.; Writing - original draft: R.S., W.H.; Writing - review & editing: R.S., F.M., M.Ts., V.S., M.Tu., W.H.; Visualization: R.S., Supervision: W.H., M.Ts., M.Tu.; Project administration: R.S., V.S.; Funding acquisition: R.S., M.Tu.

## Supplementary Figures

**Fig. S1.**
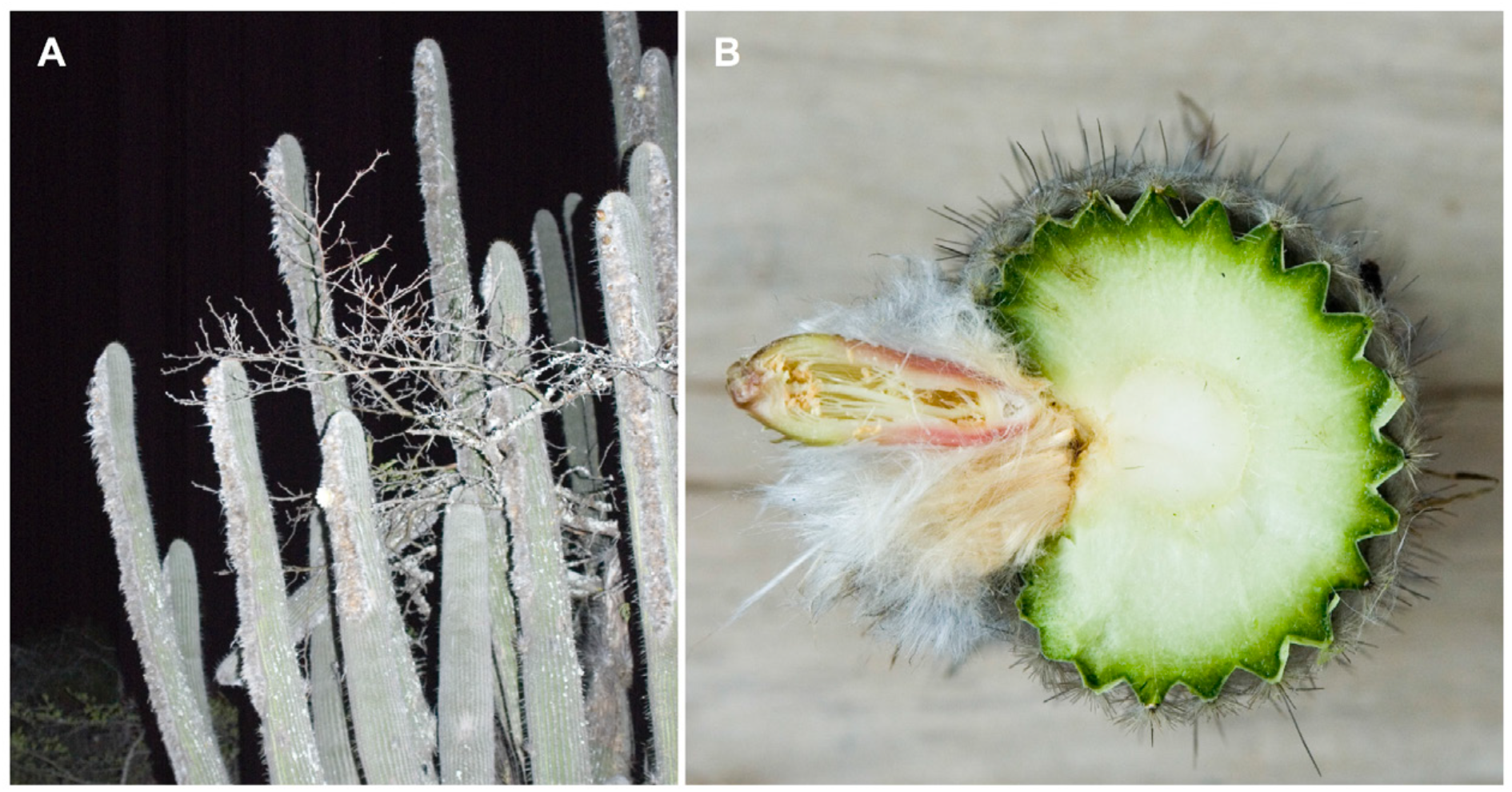
(**A**) Habitus of an *Espostoa frutescens* plant and (**B**) cross section of a column with the hairy cephalium and a closed flower.

**Fig. S2.**
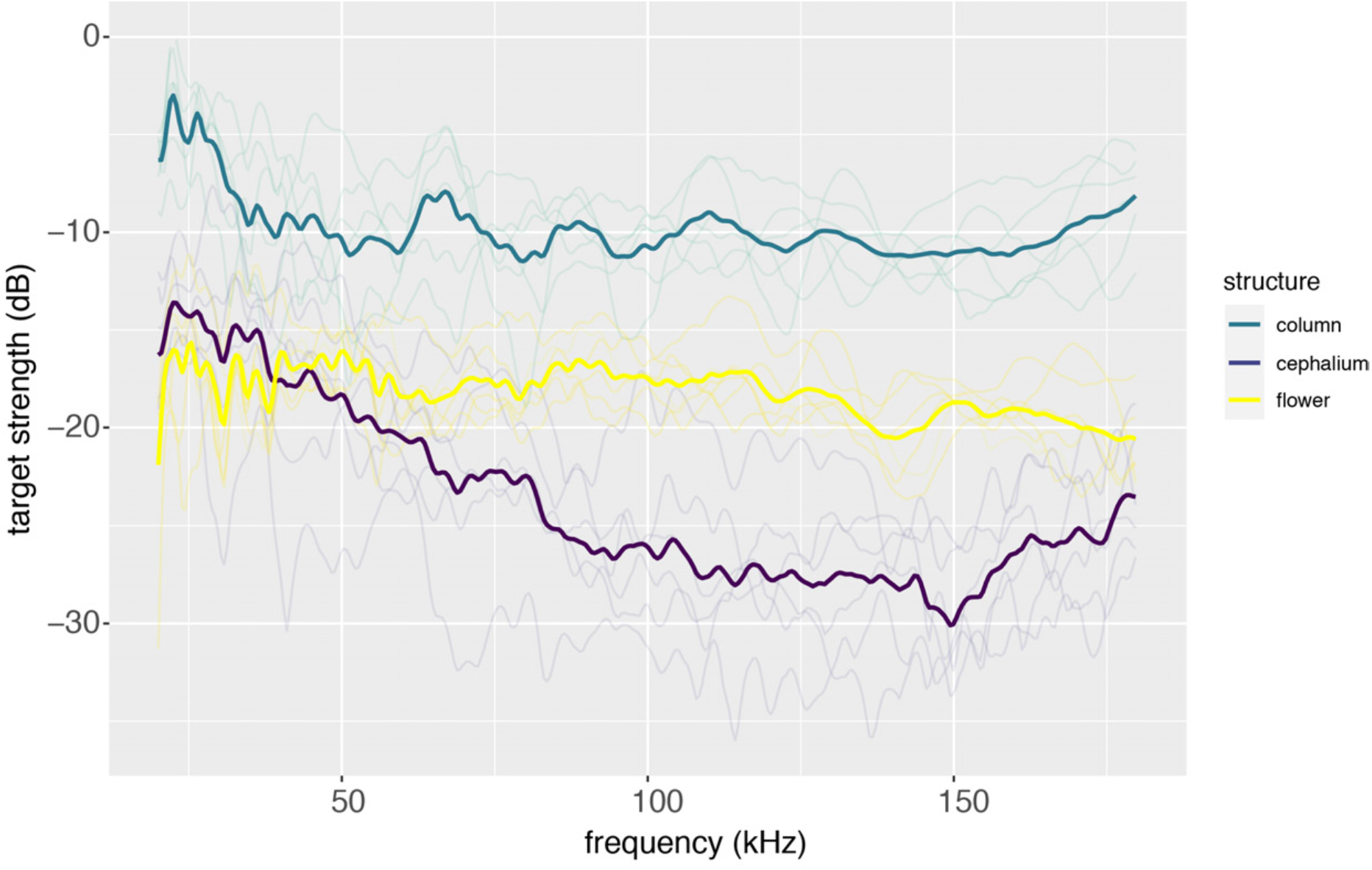
Averaged echo spectra for the three different cactus structures, column, cephalium and flower. Each single spectrum (thin, transparent lines) represents the average spectrum of 10 measurements from one cactus specimen. The bold lines give the average of 6 specimens.

**Fig. S3.**
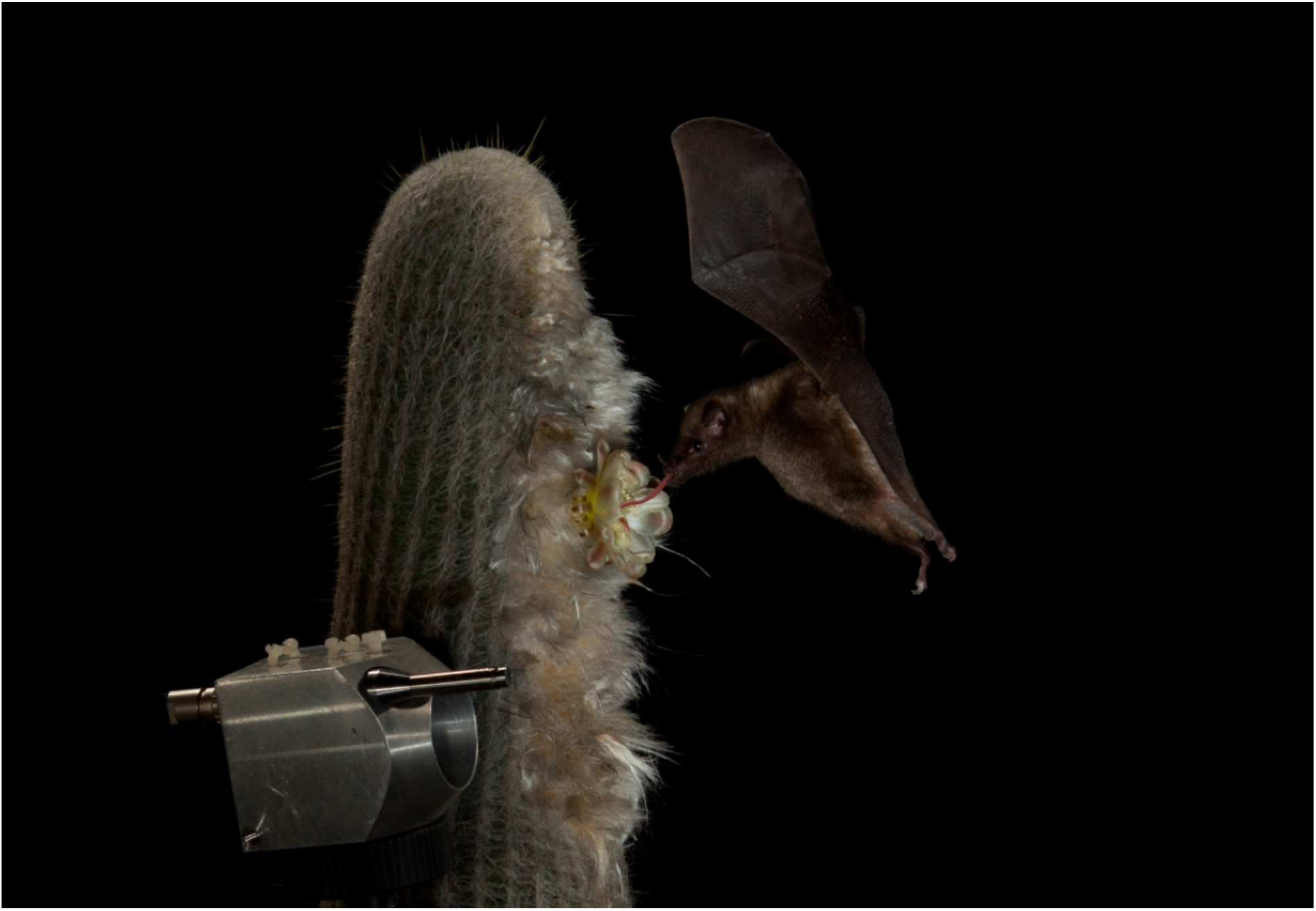
Geoffroy’s tailless bat (*Anoura geoffroyi*) drinking out of a flower of *Espostoa frutescens* while we recorded the echolocation calls during approach. We used a 1/4” G.R.A.S. microphone placed next to the flower (photo credit: Merlin Tuttle’s Bat Conservation).

